# Trimethylammonia-lyases of *Shewanella oneidensis* and Their Role in Anaerobic Respiration

**DOI:** 10.64898/2026.04.22.720169

**Authors:** Yulia V. Bertsova, Marina V. Serebryakova, Olga S. Godovanets, Victor A. Anashkin, Alexander A. Baykov, Alexander V. Bogachev

**Author notes:** Corresponding author: Belozersky Institute of Physico-Chemical Biology, Lomonosov Moscow State University, 119234 Moscow, Russia. E-mail address (A.V. Bogachev).

## Abstract

The facultatively anaerobic bacterium *Shewanella oneidensis* MR-1 contains in its genome two operons, *so_3056–3058* and *so_3299–3301*, each including genes for putative periplasmic flavocytochrome *c* and ammonia-lyase of aromatic amino acids. To determine their role in anaerobic respiration, we produced the encoded ammonia-lyases SO_3057 and SO_3299 in *Escherichia coli* and determined their substrate specificities. SO_3057 was found to cleave trimethylammonium group from ergothioneine to yield thiourocanic acid, whereas SO_3299 catalyzed a similar conversion of N(π)-methyl histidine betaine to yield N(π)-methyl urocanate. The catalytic efficiencies (*k*_cat_/*K*_m_ values) were (3–4) × 10^6^ M^−1^ s^−1^, and the pH optima of activity were between 8 and 9. Ergothioneine induced SO_3057 synthesis in anaerobic *S. oneidensis* cells and their growth, and thiourocanate stimulated respiration as an alternative terminal electron acceptor. The predicted 3D structures of the genetically coupled flavocytochromes *c* (SO_3056/58 and SO_3300/3301) are consistent with their use of thiourocanate and N(π)-methyl urocanate, respectively, as electron acceptors. We therefore conclude that the periplasmic lyases encoded by the *so_3057* and *so_3299* genes contribute to anaerobic respiration in *S. oneidensis* by producing terminal electron acceptors for the genetically coupled flavocytochromes *c*.

## 1. Introduction

In the absence of O_2_, many heterotrophs utilize various inorganic and organic compounds as terminal electron acceptors for the respiratory chain to increase the energy yield of fermentation or allow growth on various non-fermentable substrates [1-6]. While less favorable in terms of redox potential and, hence, energy yield, organic acceptors are more readily available, as they are often the products of central metabolism or simple transformations of common secondary metabolites. For example, fumarate can be synthesized from malate, aspartate, or tartrate [7]; acrylate from the osmolyte dimethylsulfoniopropionate [5, 8]; and urocanate from histidine or histidine betaine [4, 9].

The facultatively anaerobic bacterium *Shewanella oneidensis* MR-1 can use an exceptionally wide range of terminal electron acceptors (Mn(III), Mn(IV), Fe(III), Cr(VI), U(VI), fumarate, urocanate, nitrate, nitrite, trimethylamine N-oxide, dimethyl sulfoxide, sulfite, thiosulfate, and elemental sulfur) [4, 10, 11], making this bacterium a classic model for studying anaerobic respiration. Anaerobic reduction of organic acceptors is generally carried out in *Shewanella* by so-called flavocytochromes [12-14]. These proteins are localized in the periplasmic space and consist of two domains or subunits. The FAD-containing domain/subunit (Pfam ID: PF00890) is responsible for organic acceptor binding and reduction, while the heme C-containing domain (multiheme cytochrome *c*) or the domain containing covalently bound FMN (PF04205) is responsible for electron transfer from the donor of reducing equivalents to the FAD-containing domain. The donor of reducing equivalents for flavocytochromes is usually a low-potential cytochrome *c* of the respiratory chain. The genome of *S. oneidensis* MR-1 [11] contains seven sets of genes for various flavocytochromes: *so_0970, so_1413/1414, so_1421, so_3056/3058, so_3300/3301, so_3623/3624*, and *so_4620*, of which *so_3624* is possibly a pseudogene [15]. Of these, only two flavocytochromes (fumarate reductase SO_0970 [2, 12] and urocanate reductase SO_4620 [4]) have been characterized in terms of their role in anaerobic respiration and substrate specificity. It is thus plausible that the other flavocytochromes allow *S. oneidensis* to use yet unidentified organic electron acceptors for anaerobic growth.

The genes of two flavocytochromes (*so_3056/3058* and *so_3300/3301*) are located on the *S. oneidensis* chromosome within operons that additionally include genes for putative periplasmic ammonia-lyases of aromatic amino acids, *so_3057* and *so_3299* (Fig. 1A). These enzymes (PF00221) are usually homotetrameric proteins that catalyze the cleavage of the α-amino group from histidine, phenylalanine, or tyrosine to form urocanic, cinnamic, or *p*-coumaric acid, respectively [16]. The ammonia-lyases contain a unique electrophilic prosthetic group, 4-methylideneimidazole-5-one (MIO), which is formed autocatalytically from a conserved Ala-Ser-Gly sequence [17]. The MIO group is essential for catalysis and participates in substrate α-amino group removal [18], which is accompanied by a β-carbon proton transfer to a conserved Tyr residue of the ammonia-lyase, yielding a *trans* isomer of the corresponding α,β-unsaturated carboxylic acid [16]. Since this product can act as terminal electron acceptor [3], the gene clusters shown in Fig. 1A may encode alternative pathways for terminal electron acceptor formation in anaerobic respiration [11].

**Fig. 1.**
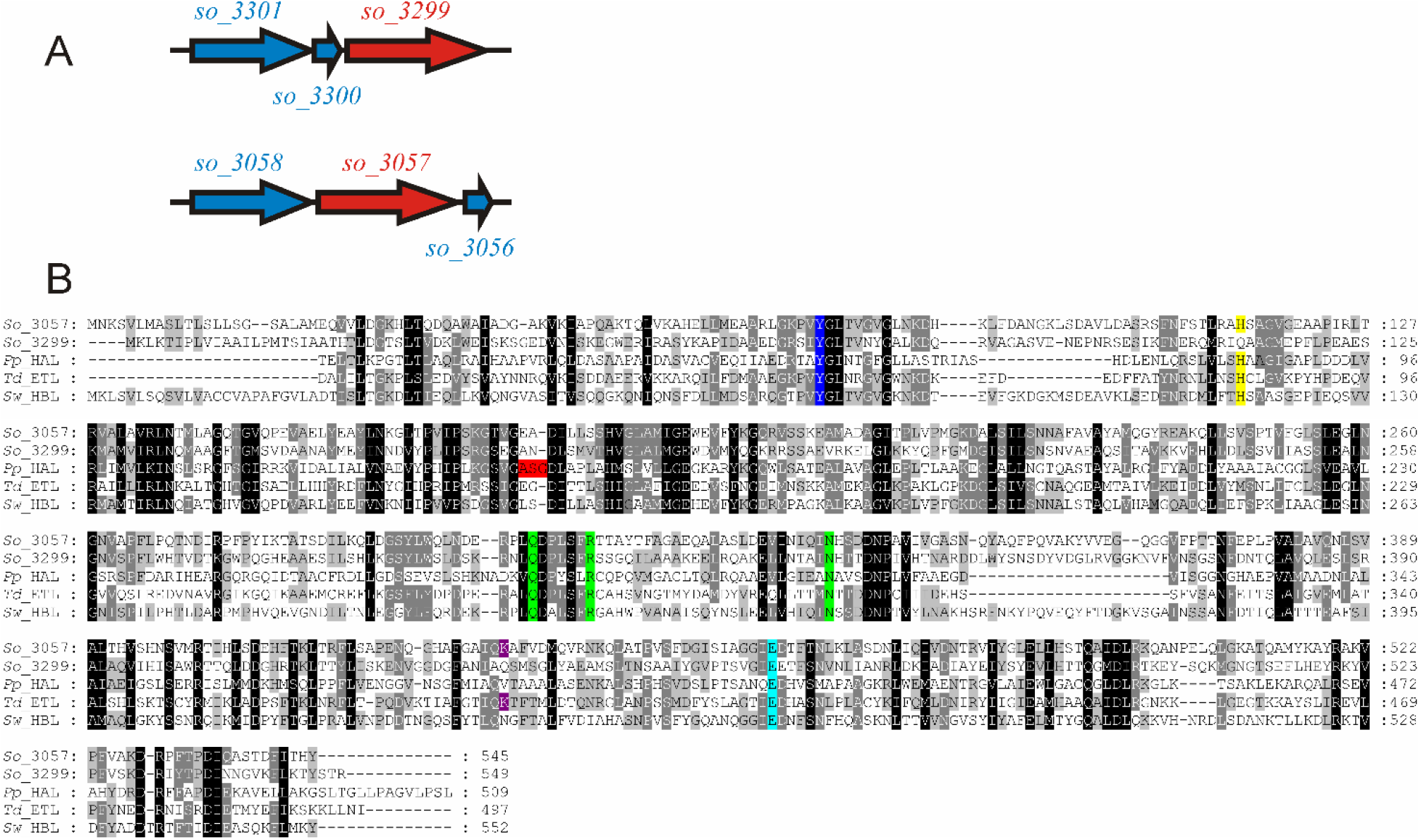
The proteins of interest encoded by *S. oneidensis* genome. (A) The gene clusters *so_3056*– *so_3058* and *so_3299*–*so_3301* in *S. oneidensis*. The genes encoding FAD- and heme C-containing subunits of flavocytochromes *c* are marked in blue, and the genes encoding putative ammonia-lyases are marked in red. (B) Multiple sequence alignment of the *S. oneidensis* SO_3057 and SO_3299 proteins, *Pseudomonas putida* histidine ammonia-lyase (*Pp*_HAL, GenBank Id: BAN56952), *T. denticola* ergothionase (*Td*_ETL, GenBank Id: EMB27117), and *S. woodyi* histidine betainase Swoo_3913 (*Sw*_HBL, GenBank Id: ACA88171). The alignment was produced with ClustalW. The residues involved in the binding of the substrate carboxylic group, imidazole N(π)-atom, and imidazole N(τ)-atom are highlighted in green, turquoise, and yellow, respectively. Other colors: blue, the Tyr residues essential for substrate β-proton abstraction; red, the Ala-Ser-Gly sequence, forming the MIO-group of ammonia-lyases; magenta, the Lys384 residue essential for ergothioneine thiolate group binding in *Td_*ETL.

The amino acid sequences of the putative lyases SO_3057 and SO_3299 (Fig. 1B) are similar to those of histidine ammonia-lyases. However, SO_3057 and SO_3299 differ from the latter enzymes in lacking the conserved Ala-Ser-Gly sequence, required for MIO group formation in all so far characterized ammonia-lyases of aromatic amino acids. MIO absence is characteristic of trimethylammonia-lyases (ergothionases and histidine betaine lyases), which cleave the trimethylammonium group from ergothioneine (Erg) or histidine betaine (HB) (Fig. 2) [9, 19-21]. SO_3057 contains all the amino acid residues necessary for ergothionase activity [20]. These include the residues involved in substrate carboxyl group, imidazole nitrogen atoms, and thiolate group binding as well as the Tyr residue, which accepts the proton from the β-carbon atom of Erg (Fig. 1B). Based on indirect evidence, Baran *et al*. suggested that SO_3057 has ergothionase and histidine betaine lyase activities [22]. However, the substrate specificity, catalytic characteristics, and physiological role of this protein in *S. oneidensis* cells remain largely unknown.

**Fig. 2.**
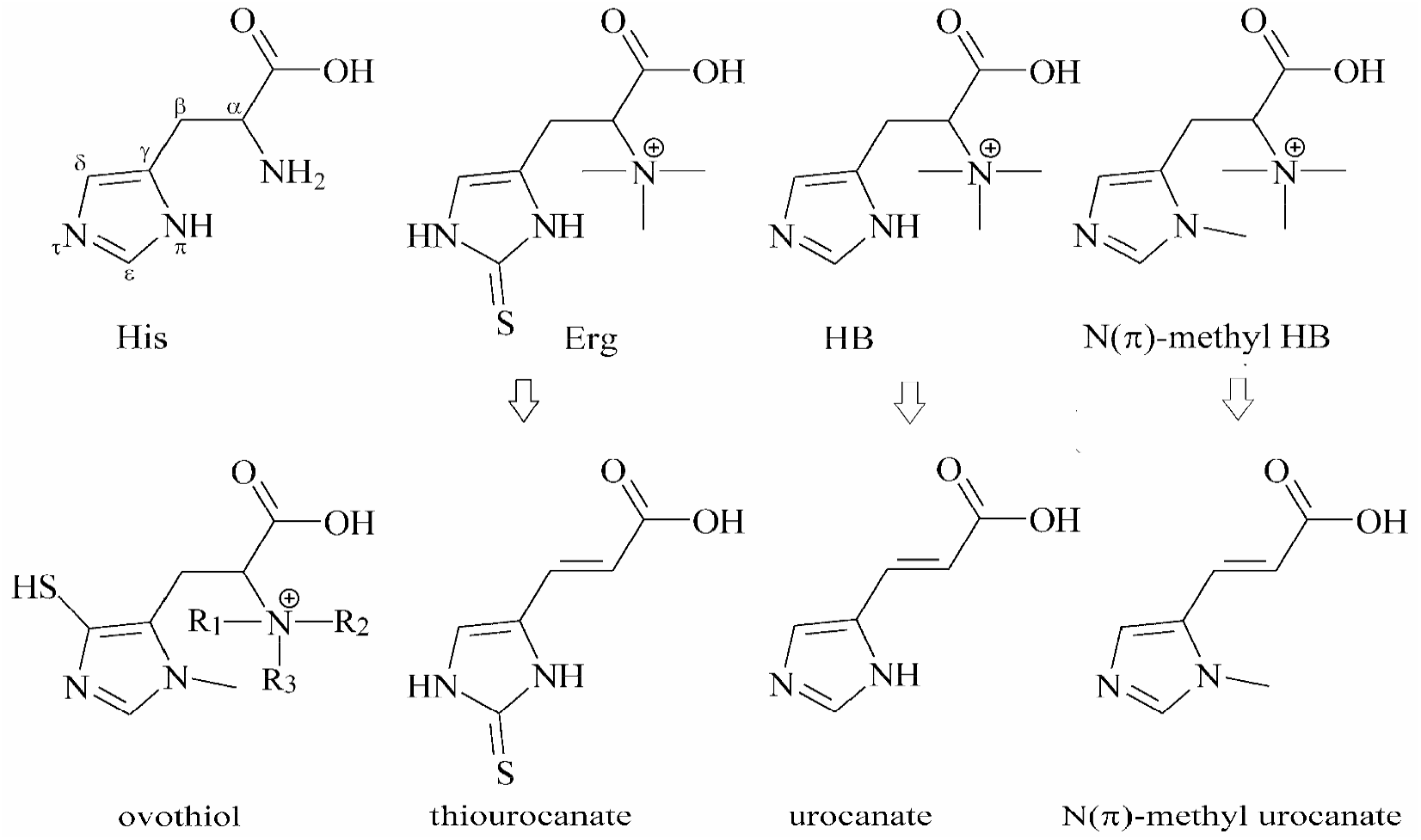
L-Histidine and its derivatives and their interconversions considered in this study. The histidine atoms are numbered according to IUPAC nomenclature. Ovothiol structure refers to three compounds: ovothiol A, R_1_ = R_2_ = R_3_ = H; ovothiol B, R_1_ = R_2_ = H, R_3_ = methyl; ovothiol C, R_1_ = H, R_2_ = R_3_ = methyl. All other substituents at the N atoms are methyl groups.

The SO_3299 protein also contains most of the amino acid residues necessary for ergothionase activity (Fig. 1B). However, SO_3299 has a neutral Gln434 residue in the position of Lys384 of *Treponema denticola* ergothionase. Notably, the Lys residue is required for the formation of the thiolate form of ergothioneine and thiolate group binding by ergothionases, and its replacement led to a loss of ergothionase activity [20]. Clearly, the SO_3299 substrate might be an ergothioneine derivative that does not contain a sulfur atom, i.e., HB [9].

This study focused on the two key enzymes encoded by the gene clusters *so_3056–so_3058* and *so_3299–so_3301*, the trimethylammonia-lyases SO_3057 and SO_3299. We heterologously produced these enzymes and determined their catalytic characteristics, including substrate specificity. We also present indirect evidence that these lyases produce terminal electron acceptors for genetically coupled flavocytochromes *c*, thereby supporting anaerobic respiration in *S. oneidensis*.

## 2. Materials and Methods

### 2.1. Materials

Ergothioneine, L-histidine betaine (HB), N(τ)-methyl-L-histidine, and N(π)-methyl-L-histidine were obtained from Shanghai Macklin Biochemical Technology Co., Shanghai, PRC. The identity and high purity (>99 %) of L-histidine betaine were confirmed by mass-spectral analysis. O-Methyl L-histidine was from Sigma Chemical Co., and L-histidine was a Reanal (Hungary) product.

Methylation of L-histidine, HB, O-methyl-L-histidine, N(τ)-methyl-L-histidine, and N(π)-methyl-L-histidine with iodomethane was performed as described for HB synthesis [23]. Histidine or its derivative (1.48 mmol) was dissolved in 3 mL of a water/methanol (2:1, v/v) mixture. The pH value was adjusted to 11 with 3 M NaOH, and 5.18 mmol of iodomethane were added with stirring. The resulting mixture was stirred at room temperature while maintaining pH 10–11 with portions of 3 M NaOH until the pH stabilized (∼6 h). Finally, the mixture was neutralized with hydrochloric acid, assayed for the presence of SO_3299 substrate(s), and subjected to mass-spectral analysis.

The product of N(π)-methyl-histidine methylation was purified using chromatography on a 1.6 × 15-cm column of Dowex 50-X4 resin (H^+^ form). After loading of the product, the column was washed with water until the pH of the eluate was approximately 7, and the methylated histidine derivative was eluted with 2 M HCl solution. Aliquots of the eluate were tested for the presence of substrate for SO_3299. The positive fractions were combined and evaporated to dryness under reduced pressure over KOH flakes. The residue was extracted with hot methanol, and crystallized by the addition of diethyl ether. The crystallization procedure was repeated three times.

### 2.2. Bacterial strains and growth conditions

*S. oneidensis* MR-1 cells were grown aerobically or anaerobically at 28°C in a medium containing 0.225 g/L K_2_HPO_4_, 0.225 g/L KH_2_PO_4_, 0.46 g/L NaCl, 0.225 g/L (NH_4_)_2_SO_4_, 0.117 g/L MgSO_4_×7H_2_O, 20 mM L-lactate, 0.05% yeast extract, and 0.1 M HEPES/NaOH (pH 7.2). Anaerobic cultivations were carried out in sealed glass flasks completely filled with the medium.

### 2.3. Genetic manipulations

To construct the plasmid encoding a cytoplasmic 6×His-tagged variant of SO_3057, the nucleotides 64–1635 of the *so_3057* gene, excluding those for N-terminal leader peptide, were amplified from the genomic DNA of *S. oneidensis* MR-1 by PCR using high fidelity Tersus polymerase (Evrogen) and the forward/reverse primers 5’-<u>CATATG</u>GAACAAGTGGTGCTT / 5’-<u>CTCGAG</u>ATAATGTGTAATAAAGTCAGT (restriction sites for *Nde*I and *Xho*I are underlined). The amplified 1581-bp fragment was cloned into a pCR4-TOPO vector (Invitrogen), resulting in a pCR_c3057 plasmid. Subsequently, we subcloned *so_3057* into the pSCodon vector (Delphi Genetics) using the *Nde*I/*Xho*I sites, resulting in the plasmid pSC_c3057. DNA sequencing was used to verify the SO_3057-encoding region of pSC_c3057, and the plasmid was transformed into the *Escherichia coli* BL21 (DE3) strain.

To construct the plasmid encoding a cytoplasmic 8×His-tagged variant of SO_3299, the nucleotides 61–1647 of the *so_3299* gene, excluding those encoding the N-terminal leader peptide, were amplified from the genomic DNA of *S. oneidensis* MR-1 by PCR using Tersus polymerase and the forward/reverse primers 5’-<u>CATATG</u>GCAACTCATACGCTTGATGG / 5’-<u>GCGGCCGC</u>TCTTGTACTGTAGGTTTTTAAAAAT (restriction sites for *Nde*I and *Not*I are underlined). The amplified 1601-bp fragment was cloned into a pAL2-T vector (Evrogen), resulting in a pAL_c3299 plasmid. Subsequently, we subcloned *so_3299* into the pET36b vector (Novagen) using the *Nde*I/*Not*I sites, resulting in the plasmid pET36_c3299. The SO_3299-encoding region of pET36_c3299 was verified by DNA sequencing, and the plasmid was transformed into *E. coli* BL21 (DE3).

### 2.4. Isolation of SO_3057 and SO_3299

Recombinant 6×His-tagged SO_3057 and SO_3299 proteins were heterologously produced in *E. coli* cells and purified by metal chelate chromatography as previously described for the *Shewanella woodyi* Swoo_3913 protein [9]. Protein concentration was determined by the bicinchoninic acid method [24] using bovine serum albumin as a standard.

### 2.5. Determination of enzymatic activities in vitro

Ergothionase, histidine betainase, and N(π)-methyl-histidine betainase activities of SO_3057 and SO_3299 were determined at 25°C by following (with an Aminco DW-2000 spectrophotometer) the formation of thiourocanate (*ε* = 22.5 mM^-1^ cm^-1^ at 311 nm), urocanate (*ε* = 18.8 mM^-1^ cm^-1^ at 277 nm), or N(π)-methyl-urocanate (*ε* ≈ 20 mM^-1^ cm^-1^at 289 nm), respectively. The assay mixture contained 0.01–2 mM substrate, SO_3057 or SO_3299 (0.03–10 μg/mL), and 50 mM Bis-Tris propane-HCl buffer (pH 8.5).

The pH-dependence of the enzymatic activities (reaction rates *v*) was measured with 0.5 mM substrate in a series of 0.1 M Bis-Tris propane buffers adjusted to appropriate pH values with citric acid. The data were treated using the following equation:

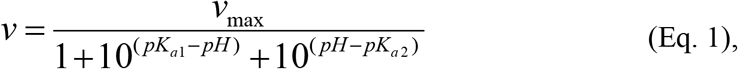

where *v*_max_ is the theoretical limiting value of the rate *v* and p*K*_a1_ and p*K*_a2_ are the acid dissociation constants for two catalytically important groups.

### 2.6. Determination of ergothionase activity in S. oneidensis cells

*S. oneidensis* cells were grown aerobically or anaerobically with an appropriate electron acceptor, harvested by centrifugation (11,000*g*, 2 min), and washed twice with a medium containing 85 mM NaCl, 5 mM MgSO_4_, and 10 mM Tris-HCl (pH 8.0). The cells were added to the assay medium containing 0.1 M Tris-HCl (pH 8.5) and 0.5 mM Erg, and the enzymatic reaction was followed spectrophotometrically at 311 nm (*ε* = 22.5 mM^-1^ cm^-1^).

### 2.7. Mass-spectral analyses

The procedures for MALDI-TOF MS analysis of the purified proteins have been previously described [25]. The proteins were separated by SDS-PAGE and in-gel digested with trypsin. Tryptic peptides were analyzed using Bruker Daltonics UltrafleXtreme MALDI-TOF-TOF mass spectrometer (Germany) equipped with a UV laser (Nd) in a positive ion mode using a reflectron. Probe solution (0.1-1 μL) was mixed on a target plate with 0.5 μL of 2,5-dihydroxybenzoic acid (Aldrich, 40 mg/mL in 30% aqueous acetonitrile, 0.5% trifluoroacetic acid). The accuracy of the measured monoisotopic masses was within 50 ppm. Protein identification was performed using peptide fingerprint option of Mascot ver. 2-3-02 (www.matrixscience.com) in a Home database, taking into account possible oxidation of methionines and possible modification of cysteines by acrylamide in the gel.

The same mass spectrometry modes were used to analyze histidine derivatives and the products of their conversion by SO_3299. To obtain the fragmentation spectra of these compounds [26], the mass spectrometer was used in a tandem mode; the accuracy of the fragment ion measurements was within 1 Da.

### 2.8. Bioinformatics

Three-dimensional structure of mature FAD-containing subunits of the *S. oneidensis* flavocytochromes SO_3058 and SO_3301 were predicted from their amino acid sequences using the program Boltz-2 (version 2.2.1) [27]. Default parameter values were used, and the numbers of recycling steps and generated diffusion samples were set to 10 and 25, respectively. Each model contained bound FAD and the substrate—N(π)-methylurocanate for SO_3301 or *ε*-thiourocanate for SO_3058. N(π)-Metylurocanate was present in its zwitterionic form with protonated N(τ) atom, and *ε*-thiouroocante as a thione. Ligand affinities were calculated for 20 diffusion samples. The models with the best confidence scores (typically 0.965−0.980) were finally selected. Ramachandran plots showed that not less than 97.7 % amino acid residues were in allowed conformations. All calculation experiments were performed in triplicate, using a graphical station with an NVIDIA RTX 4000 Ada Generation GPU accelerator.

The program PLIP was used to detect π–π interactions in protein structures [28]. Multiple sequence alignments were obtained with ClustalW (https://www.genome.jp/tools-bin/clustalw) using default parameter settings. Cellular localization of bacterial proteins was predicted using SignalP 6.0 [29]. UCSF Chimera (version 1.19) was used for structure visualization [30].

## 3. Results

### 3.1. Heterologous production of the SO_3057 and SO_3299 proteins in E. coli cells

The genes encoding the truncated SO_3057 and SO_3299 proteins, lacking signal peptides, were amplified from *S. oneidensis* genomic DNA, cloned into expression vectors, and the corresponding proteins were produced in *E. coli* cells. The recombinant SO_3057 and SO_3299 proteins were isolated in electrophoretically pure forms (Fig. S1) using metal-affinity chromatography with yields of 130 and 180 mg per 1 liter of *E. coli* culture, respectively. Mass spectrometric analysis of the bands obtained after polyacrylamide gel electrophoresis confirmed the identification of SO_3057 (sequence coverage: 100%) and SO_3299 (sequence coverage: 70%).

### 3.2. The substrate specificities of SO_3057 and SO_3299

Previous mass spectrometric studies suggested that SO_3057 is capable of cleaving the trimethylammonium group from Erg and HB, i.e., that this protein exhibits ergothionase and histidine betainase activities [22]. In accordance with this prediction, SO_3057 catalyzed Erg conversion into thiourocanate, as evidenced by the decrease in Erg absorbance at 257 nm and increase in thiourocanate absorbance at 311 nm (Fig. 3). However, in contrast to previously obtained results [22], we were unable to detect any ability of SO_3057 to cleave the trimethylamino group from the commercial high-purity HB used in this study. On the other hand, the inability of SO_3057 to convert HB is in a good agreement with the properties of other described ergothionases [20].

**Fig. 3.**
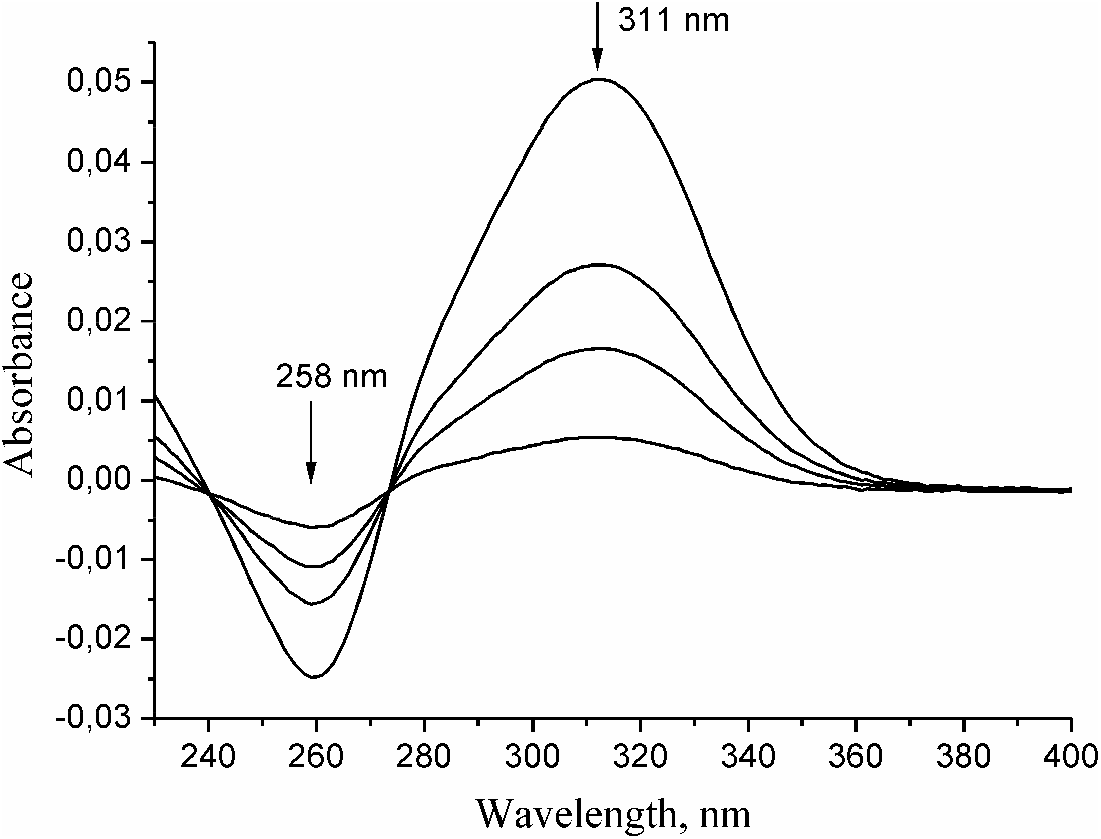
The catalytic activity of SO_3057 with ergothioneine (Erg) as substrate. Spectral changes in the assay medium during the enzymatic conversion of Erg are shown; the spectra from bottom to top were recorded after 30, 90, 180, and 360 s after adding 0.026 μg/mL SO_3057 to the assay medium containing 50 μM Erg and 0.05 M Bis-Tris propane-HCl (pH 8.5).

The other putative trimethylammonia-lyase, SO_3299, lacks the conserved Lys residue and, by analogy with the earlier characterized histidine betainase [9], was expected to convert HB but not Erg (see Introduction). This prediction was only partly confirmed—SO_3299 demonstrated activity with neither of these compounds.

Notably, the lack of HB conversion was established with the high-purity HB preparation. However, some activity was detected with the HB preparation obtained in this laboratory by histidine methylation with iodomethane (Fig. 4A). Importantly, the UV spectrum of the reaction product differed from that of urocanic acid, the expected product of HB conversion, in peak wavelength (289 nm versus the expected 277 nm [9]) (Fig. 4B). Furthermore, the reaction ceased shortly, resulting in only a small absorbance change, which could, however, be reproduced by the addition of a fresh portion of the substrate (Fig. 4A). These findings suggested that the SO_3299 substrate is a minor product of histidine methylation (∼ 2 % of the total methylated histidine content).

**Fig. 4.**
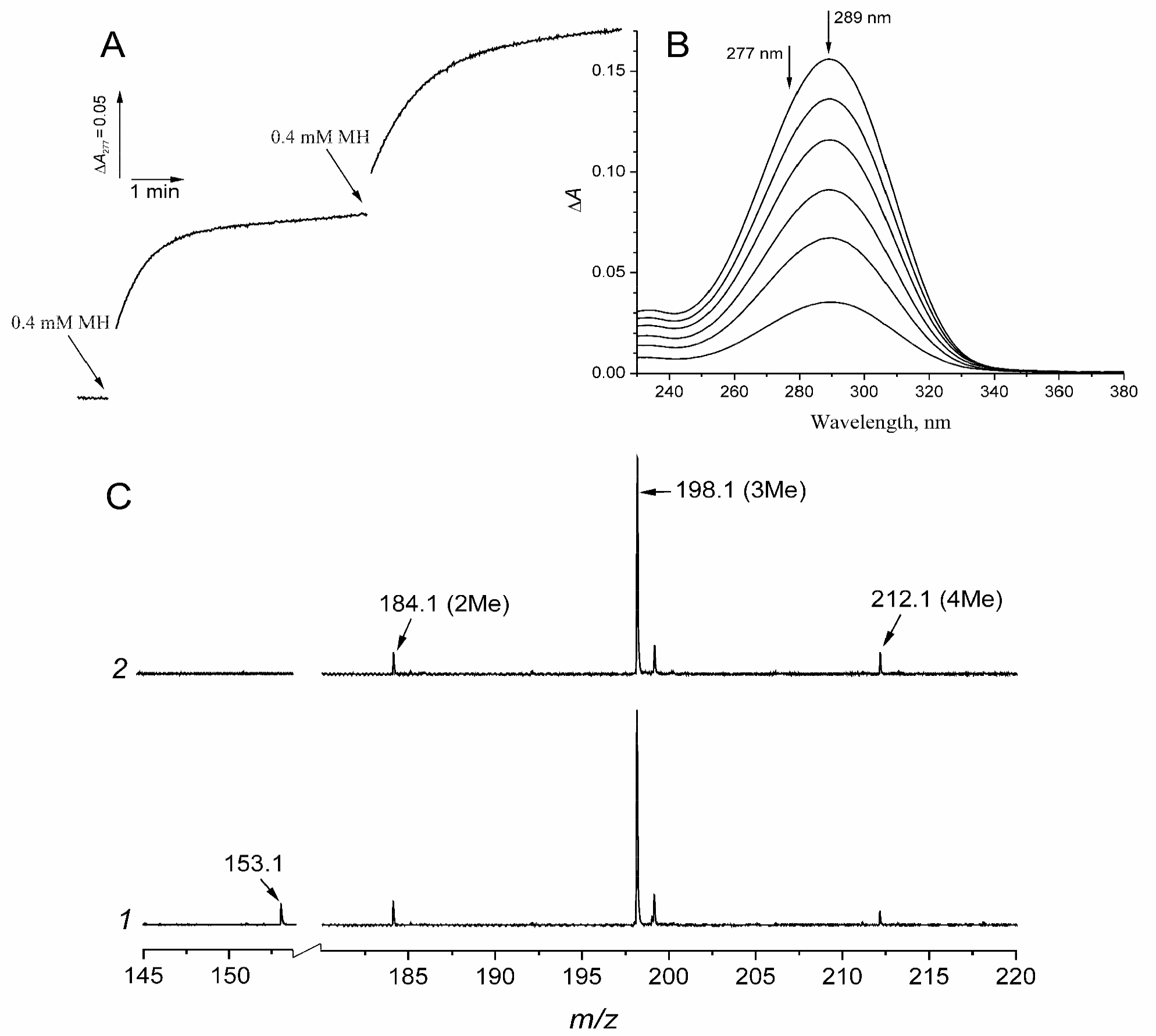
The catalytic activity of SO_3299 with methylated L-histidine as substrate. (A) The time-courses of light absorbance at 277 nm measured with 4 μg/mL SO_3299 and two portions of ∼ 0.1 mM crude methylated histidine (MH) in 0.1 M Tris-HCl buffer (pH 8.5). (B) The spectral changes in the assay medium during the enzymatic conversion of the methylated histidine preparation; the reaction time increased from 30 s (bottom) to 180 s (top) in 30-s intervals. (C) The mass spectra of the MH preparation (∼ 1 mM) incubated for 20 min with 1 μg/mL native SO_3299 (spectrum *1*) or denatured SO_3299 (spectrum *2*). The two spectra differ in only the order of SO_3299 and trifluoroacetic acid (5 %) additions to MH—the acid was added either after incubation with SO_3299 (spectrum *1*) or before it (spectrum *2*). Two regions of the spectra, containing the peaks corresponding to methylurocanate and histidine with two to four added methyl group(s) are shown. The peaks are indicated by arrows and labelled on one or the other spectrum.

Mass-spectral analysis of the methylated histidine preparation obtained by the reaction of histidine with iodomethane helped to identify the true substrate of SO_3299. The spectrum of the crude preparation (Fig. 4C) contained the peaks of His(methyl)_2_ (*m/z* = 184.1), His(methyl)_3_ (*m/z* = 198.1, matching HB), and His(methyl)_4_ (*m/z* = 212.1), with amplitude ratios of ∼0.1:1:0.1. If this preparation was incubated with SO_3299 until the reaction stopped, the His(methyl)_4_ peak decreased and an additional peak with *m/z* = 153.1 (matching methylurocanate) appeared in the products (Fig. 4C). The fact that the peak of His(methyl)_4_ did not disappear upon prolong incubation with SO_3299 (Fig. 4C) suggested that its significant fraction refers to the non-convertible species with a different distribution of four methyl groups among the possible methylation sites.

To determine the position of the only methyl group retained in the product of the enzymatic reaction, we performed methylation of histidine derivatives, each already containing a methyl or trimethyl substituent at one of the four possible locations, and tested the products as SO_3299 substrates. Notably, none of these compounds was converted by SO_3299 before this additional methylation. The conditions for the methylation reaction (starting substance:iodomethane ratio = 1:3.5, pH 10–11) were optimal for α-amino group methylation in histidine and its analogs [23]. As described above, histidine methylation yielded approximately 2 % molar yield of SO_3299 substrate. The yield increased to 7 % with methylated commercial HB preparation (Fig. 5), indicating the requirement for an α-trimethylamino group in SO_3299 substrate. The highest yield of the SO_3299 substrate (∼50%) was obtained with N(π)-methyl histidine. In contrast, methylation of N(τ)-methyl histidine and O-methyl histidine did not yield any SO_3299 substrate.

**Fig. 5.**
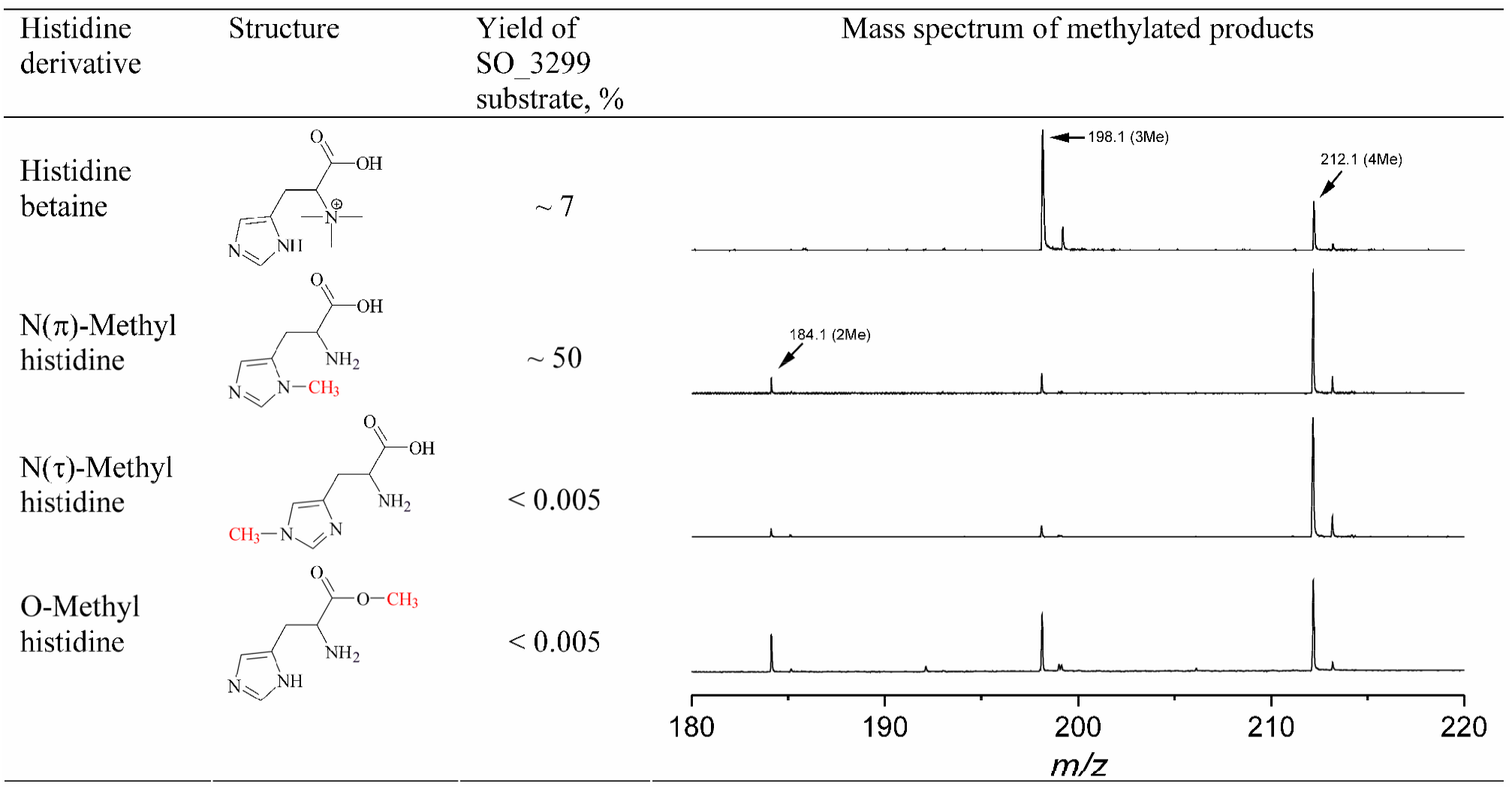
Formation of SO_3299 substrate in the reaction of various histidine derivatives with iodomethane. The yields were determined spectrophotometrically using SO_3299-catalyzed reaction to convert the substrate present among the methylated products to methylurocanic acid (*ε*_289_ ≈ 20 mM^-1^ cm^-1^). Yield was calculated in each case as the ratio of the amount of the methylurocanic acid formed in the enzymatic reaction and the amount of the histidine derivative reacted with iodomethane. No methylurocanic acid or any other product absorbing light at 230−380 nm was detected after reaction of SO_3299 with the methylated forms of two last histidine derivatives.

Mass spectral analysis of the methylated preparations (Fig. 5) identified His(CH_3_)_4_ as the major product of N(π)-methyl histidine, N(τ)-methyl histidine, and O-methyl histidine methylation. These findings ruled out the possibility that the inefficiency of the methylated forms of the last two compounds as SO_3299 substrates (Fig. 5) is explained by low yield in the methylation reaction. A more likely explanation is that the His(CH_3_)_4_ species produced from the latter compounds do not contain the indispensable N(π)-methyl group or their N(τ)- and O-methyl groups are strongly inhibitory.

An evident corollary is that the SO_3299 substrate is N(π)-methyl-histidine betaine, having a trimethylamino group in the α-position and a methyl group in the N(π)-position (Fig. 2). This conclusion is consistent with an ∼12-nm spectral bathochromic shift observed in SO_3299 product compared to urocanate (Fig. 4B), keeping in mind a similar shift in absorbance peak upon methylation of urocanic acid ester at the N(π) but not N(τ) position [31].

### 3.3. The catalytic properties of SO_3057 and SO_3299

The dependence of the initial rate of the enzymatic reaction catalyzed by SO_3057 on Erg concentration (Fig. S2A) was hyperbolic, with the following catalytic parameters: *k*_cat_ = 120 ± 4 s^−1^, *K*_m_ = 41 ± 3 μM, *k*_cat_/*K*_m_ = 2.8 × 10^6^ M^−1^ s^−1^. The catalytic efficiency of SO_3057 towards Erg was thus quite high and similar to those of previously described ergothionases from *Burkholderia sp*. HME13 and *T. denticola* [19, 20]. The pH-dependence of SO_3057 activity, measured at a saturating Erg concentration of 0.5 mM, was bell-shaped (Fig. S2B), with a pH optimum in the alkaline range (8.0–8.5) and the p*K*_a_ values of 6.5 and 10.7 for two catalytically important ionizable groups.

The dependence of SO_3299 lyase activity on N(π)-methyl-HB concentration was also hyperbolic (Fig. S3A), yielding the following approximate parameter values: *K*_m_ ∼ 40 ± 3 μM and *k*_cat_ ∼165 ± 5 s^-1^. These values are approximate as they were obtained assuming the extinction coefficient *ε*_289nm_ of 20 mM^-1^·cm^-1^ for N(π)-methylurocanate, by analogy with the values reported for urocanate (*ε*_277nm_ = 18.8 mM^-1^·cm^-1^) and thiourocanate (*ε*_311nm_ = 22.5 mM^-1^·cm^-1^). The ratio *k*_cat_/*K*_m_ (catalytic efficiency) of 4.1 × 10^6^ M^-1^·s^-1^, obtained from the above parameter values, does not depend on the accuracy of the extinction coefficient because it affects *k*_cat_ and *K*_m_ identically. The pH-dependence of SO_3299 activity, measured at a saturating N(π)-methyl-HB concentration of 0.5 mM, was bell-shaped (Fig. S3B), with a pH optimum in the alkaline range (8.2–9.2) and p*K*_a_ values of 7.0 and 10.3 for two catalytically important ionizable groups.

### 3.4. SO_3057 supports anaerobic respiration in S. oneidensis

Enzymes involved in bacterial anaerobic respiration are commonly induced under conditions favorable for using the respective electron acceptor, i.e. at O_2_ limitation and in the presence of a suitable substrate [4, 6, 25, 32]. Table 1 shows the effects of growth conditions on the level of ergothionase activity in *S. oneidensis* cells. Under aerobic conditions, the activity was quite low and was not induced by Erg, and anaerobiosis alone only slightly stimulated it. In contrast, the ergothionase activity increased dramatically (∼300-fold) by Erg in the absence of O_2_. These findings strongly supported potential SO_3057 involvement in anaerobic respiration in *S. oneidensis*.

**Table 1.** Ergothionase activity in *S. oneidensis* cells grown under different conditions.

| Growth conditions | Ergothionase activity<sup>a</sup><br>(nmol/min per 1 mg protein) |
| --- | --- |
| +O<sub>2</sub> | 8 ± 3 |
| +O<sub>2</sub> + 10 mM Erg | 8 ± 2 |
| -O<sub>2</sub> + 10 mM fumarate | 16 ± 4 |
| -O<sub>2</sub> + 10 mM fumarate +10 mM Erg | 5,100 ± 300 |

<sup>a</sup> The mean ± SD for two independent growth experiments.

To determine the ability of Erg to support anaerobic *S. oneidensis* growth, cells of this bacterium were grown anaerobically in the presence of Erg, using the classical electron acceptor fumarate as a control. No cell growth was detected in the absence of an electron acceptor in the medium (Fig. 6), whereas 10 mM fumarate induced rapid growth, in accordance with previous findings [10]. The *S. oneidensis* cells incubated in the presence of 10 mM Erg demonstrated a similar growth rate, supporting the role of Erg as electron acceptor or its precursor for anaerobic growth.

**Fig. 6.**
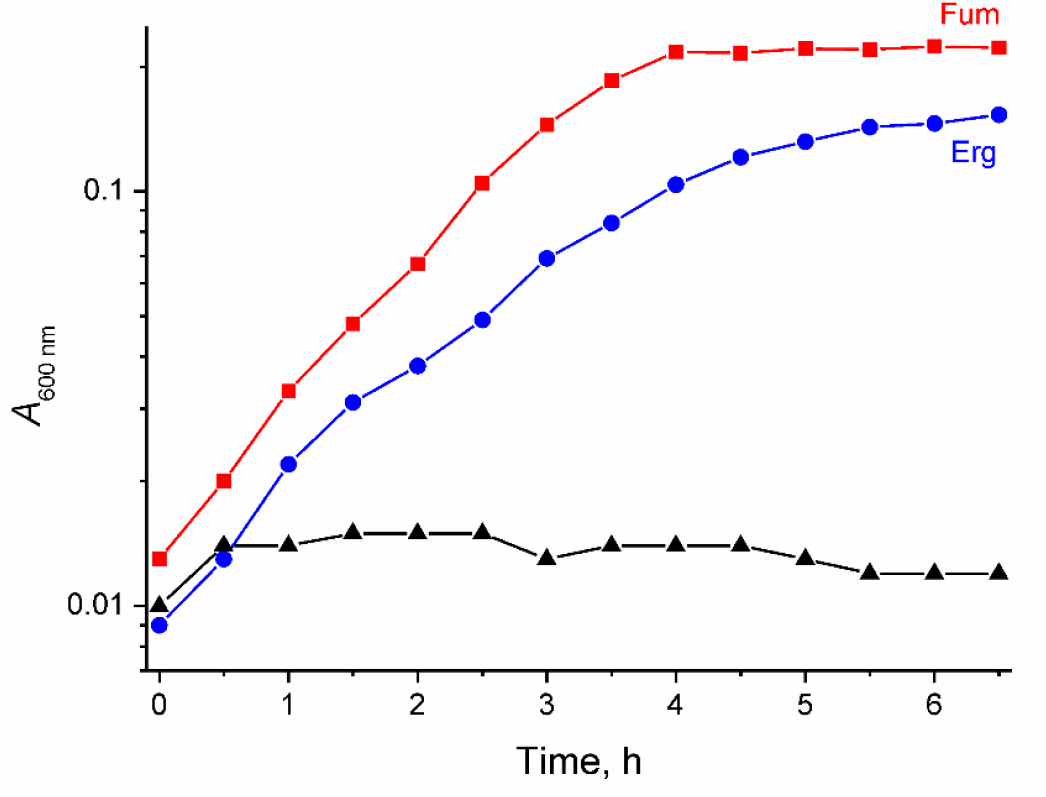
*S. oneidensis* growth curves measured in minimal medium under anaerobic conditions. Black, without additions; blue, in the presence of 10 mM ergothioneine (Erg); red, in the presence of 10 mM fumarate (Fum).

The above findings raised the possibility that *S. oneidensis* cells can use Erg or the product of its conversion by SO_3057, thiourocanate, as an alternative terminal electron acceptor. The latter alternative was tested by determining the ability of the induced (grown anaerobically in the presence of Erg) *S. oneidensis* cells to produce thiourocanate. As the left panel of Fig. 7 highlights, Erg addition to cells under aerobic conditions rapidly resulted in thiourocanate accumulation. Notably, the thiourocanate yield decreased markedly under anaerobic conditions in the presence of lactate, a potential electron donor for the respiratory chain. A likely explanation was that anaerobic *S. oneidensis* cells possess an additional enzymatic activity allowing rapid consumption of the produced thiourocanate. This explanation was supported by the observation that Erg addition to the anaerobic cells in the presence of reduced methyl viologen (MV) caused rapid oxidation of this electron donor (Fig. 7, right panel). This effect was not observed with the aerobically grown *S. oneidensis* cells (data not shown). Together, these findings indicated that the induced *S. oneidensis* cells can efficiently use Erg-derived thiourocanate as terminal electron acceptor in the anaerobic respiratory chain.

**Fig. 7.**
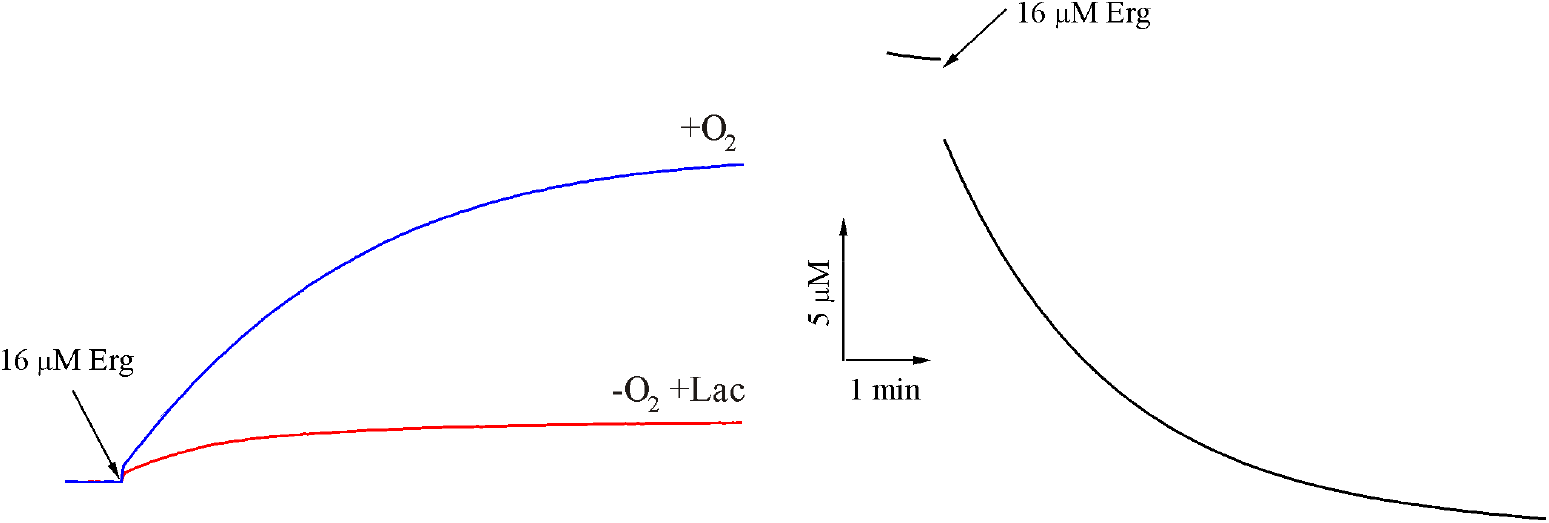
Thiourocanate formation (left) and MV oxidation (right) by *S. oneidensis* cells pre-grown anaerobically in the presence of Erg. Both reactions were followed spectrophotometrically at 311 or 606 nm, respectively. The curves were normalized by assuming that formation of one molecule of urocanate requires oxidation of two molecules of MV. The medium, used in the aerobic experiment (blue curve), contained *S. oneidensis* cells (5 μg protein/mL), 16 µM Erg, and 0.1 M Tris-HCl (pH 8.5). For anaerobic experiments, this basic medium was supplemented with 5 U/mL glucose oxidase, 5 U/mL catalase, 10 mM glucose, and 2 mM L-lactate (Lac) (red curve) or 100 μM reduced MV (black curve). The reactions were started by adding Erg.

### 3.5. Predicting substrate specificities of the flavocytochromes SO_3056/3058 and SO_3300/3301

The results reported above suggested that the flavocytochromes SO_3056/3058 and SO_3300/3301 function as thiourocanate and N(π)-methylurocanate reductases, respectively. Obtaining recombinant forms of these proteins to directly test these hypotheses was complicated by a requirement for the cell machinery allowing specific covalent bonding of heme C. Therefore, we approached the problem indirectly, by modelling the three-dimensional structures of the putative reductases with the program Boltz-2 [27] and comparing them with that of the previously characterized urocanate reductase UrdA (SO_4620). Importantly, Boltz-2 allows modelling the structures of proteins in complexes with substrates and other ligands and estimating the stabilities of such complexes.

The reference structure of UrdA (PDB ID: 6T87) contains the substrate urocanate [33] in close proximity to FAD (Fig. 8A). This positioning allows rapid hydride ion transfer from the flavin N5 atom to the substrate C3 atom and proton transfer from Arg411 to the urocanate C2 atom [33, 34]. Substrate carboxylate binding involves three residues, Arg560, Arg411, and His520 that are conserved in all 2-enoate reductases (Fig. S4) [14]. The bound urocanate molecule adopts a twisted conformation, not observed in the free state. In this conformation, the imidazole ring is rotated out of the carboxylate plane due to interaction with Glu177 and Asp388 carboxylates, which disallows π–π interactions with Tyr373 [33, 34]. The presence of negatively charged residues in the positions corresponding to UrdA Asp388 and Glu177 appears to be typical for urocanate reductases and distinguishes them from other 2-enoate reductases [14].

**Fig. 8.**
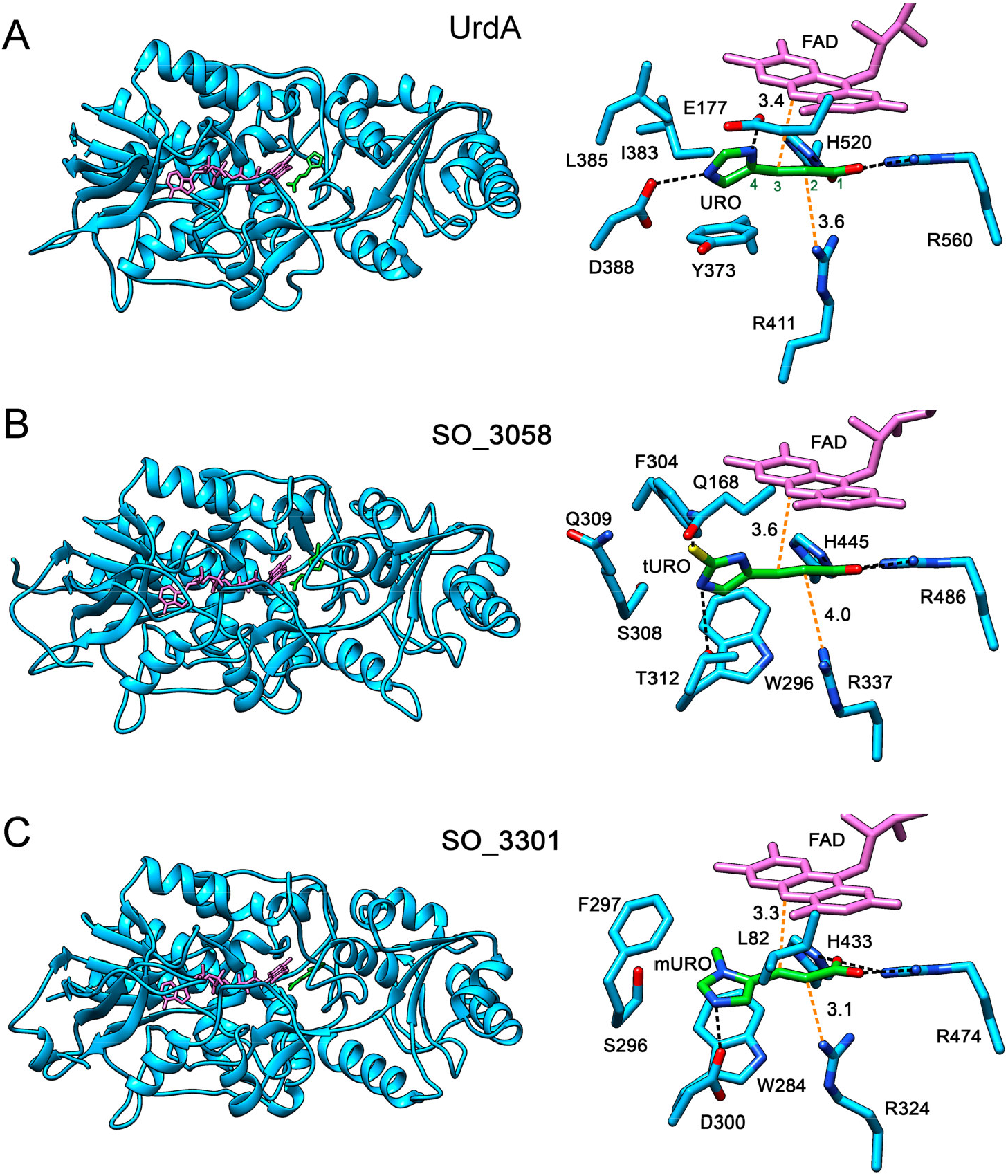
Three-dimensional structures of the flavocytochromes complexed with FAD and respective substrate. (A) The crystal structure of the urocanate reductase UrdA (SO_4620) from *S. oneidensis* with bound urocanate (URO) (PDB ID: 6T87 [33]). The C atoms of urocanate are numbered in green. (B) Predicted structure of SO_3058 with bound thiourocanate (tURO). (C) Predicted structure of SO_3301 with bound N(π)-methylurocanate (mURO). The right panels present an enlarged view of the active site in a different projection. The polypeptides are shown in turquoise, FAD in red, substrate (carbon skeleton) in green. Black dashed lines show hydrogen bonds, red dashed lines indicate the distance (shown in Å) between the FAD N5 and substrate C3 atoms and between the Arg N(ω) and substrate C2 atoms.

The modelled structures of SO_3058 and SO_3301 were similar to that of UrdA (r.m.s.d. values of 3.0 and 3.1 Å, respectively) (Figs 8B and 8C). Both modelled structures retained the cofactor FAD and substrates thiourocanate and N(π)-methylurocanate at the same sites as observed for urocanate in the UrdA structure. The binding constants for thiourocanate and N(π)-methylurocanate estimated with Boltz-2 were 35 ± 1 and 92 ± 2 µM (Δ*G* = -6.1 ± 0.2 and -5.5 ± 0.1 kcal/mol), respectively. However, the binding poses for the substituted urocanates were somewhat different. Whereas the carboxylic group of all the three substrates formed nearly identical interactions with the Arg/Arg/His triplet, the interactions and orientations of the imidazole group varied. Thiourocanate similarly bound to SO_3058 in a twisted conformation stabilized by interactions with two protein ligands but of a different nature—Gln168 and Thr312 (Fig. 8B) instead of Asp177 and Glu388 in UrdA (Fig. 8A). Importantly, the respective Gln and Thr residues are highly conserved in the SO_3058 homologs (Fig. S5) belonging to the operons additionally containing an ergothionase gene. Furthermore, the program PLIP detected two π–π interactions of the substrate imidazole group with Phe304 and Trp296, whose aromatic groups are perpendicular to the imidazole plane. Notably, the absence of negatively charged ligands for the imidazole N atoms in SO_3058 active site suggests that its substrate (thiourocanate) is bound in a thione form.

In contrast, the imidazole group of N(π)-methylurocanate participated in only one ionic interaction, with Asp300 through the N(τ) atom, and adopted conformation, permitting its stacking with Trp284 (Fig. 8C). A similar H-bond with the imidazole N(π) atom is not formed because it is bonded to a methyl group in this substrate and the UrdA N(π) ligand Asp177 is replaced by Leu in SO_3301 (Fig. S4). Methylation of the imidazole nitrogen atom likely stabilizes a particular tautomeric form of (π)-methyl urocanate with a positive charge localized on the N(τ) atom.

Importantly, the distances between thiourocanate C3 and FAD C2 atoms (3.6 Å) and between thiourocanate C2 and Arg337 N(ω) atoms (4.0 Å) in SO_3058 are short enough to allow rapid transfer of hydride ion and proton, respectively. The distances are even shorter in SO_3301 (Fig. 8C). Altogether, the modelled structures of the complexes of the flavocytochromes SO_3058 and SO_3301 with their presumed substrates provide strong support for their use of thiourocanate and N(π)-methyl urocanate as terminal electron acceptors.

## 4. Discussion

The results reported above characterize the substrate specificities and kinetic properties of two trimethylammonia-lyases (SO_3057 and SO_3299) from the facultatively anaerobic bacterium *S. oneidensis*. It was previously claimed, based on indirect evidence, that SO_3057 possesses ergothionase and histidine betainase activities [22]. Our data indeed demonstrated that this protein catalyzes trimethylammonium group cleavage from ergothioneine to yield thiourocanic acid (Fig. 3) with a catalytic efficiency *k*_cat_/*K*_m_ of 2.8 × 10^6^ M^-1^·s^-1^, similar to those of other characterized bacterial ergothionases [19, 20]. However, we did not detect any histidine betainase activity of SO_3057, which disagrees with the earlier prediction [22], while being in full concordance with the properties of other described ergothionases [20].

Typically, ergothionases have a cytoplasmic localization in bacterial cells and catalyze the first step of the aerobic Erg degradation pathway [35]. However, the SO_3057 sequence contains a leader peptide directing the protein to the periplasm. Furthermore, *so_3057* is located between two genes encoding FAD- and heme C-containing subunits of a periplasmic flavocytochrome *c* (Fig. 1A) within the same operon. In this context, it is logical to assume that the thiourocanic acid produced by ergothionase activity serves as a substrate for periplasmic flavocytochrome *c*, enabling anaerobic respiration using thiourocanic acid as a terminal electron acceptor. Notably, ergothioneine, a natural antioxidant, is widespread in the biosphere [36] and likely available to *S. oneidensis* in sufficient quantities to support anaerobic respiration.

Other anaerobic bacteria, such as *Blautia producta* and *Clostridium symbiosum*, are also known to use Erg as a precursor of the terminal electron acceptor thiourocanic acid [37]. However, its electron-accepting function is realized via a reductive desulfurization reaction catalyzed by thiourocanate desulfhydrase. *S. oneidensis* apparently employs a different strategy and uses, by analogy with *S. woodyi* urocanate reductase [9], flavocytochrome *c* to reduce the α,β-double bond of thiourocanic acid. Experimental verification of the substrate specificity of the flavocytochrome SO_3056/3058 is hindered by the inability to produce it heterologously. However, the substrate preference could be predicted from the modelled three-dimensional structure of this respiratory enzyme. According to the structure (Fig. 8B), the active site of SO_3058 is of sufficient size and possesses the residues necessary for thiourocanate binding to allow hydride ion transfer from FAD and proton transfer from the primary donor (Arg337) to the α,β-double bond. Furthermore, the active site structure is tuned to bind the thione form of thiourocanate due to absence of anionic ligands for imidazole N-atoms and a suitable positioning of the Gln168 residue that interacts with the sulfur atom and is likely the key determinant of flavocytochrome specificity towards thiourocanic acid.

Consistent with the predicted physiological role of SO_3057, high ergothionase activity was observed in *S. oneidensis* cells only when grown under anaerobic conditions in the presence of Erg (Table 1). Moreover, Erg addition to induced cells under anaerobic conditions led to respiratory activity (Fig. 7), and the presence of Erg in the medium conferred to *S. oneidensis* cells the ability to grow anaerobically (Fig. 6) with growth characteristics comparable to those using a classical electron acceptor (fumarate).

In addition to the ergothionase SO_3057, the *S. oneidensis* genome encodes another periplasmic trimethylammonia-lyase, SO_3299. Again, the operon containing its gene additionally contains the genes *so_3300–3301* for a different flavocytochrome *c* (Fig. 1A). According to our findings, SO_3299 is unable to cleave Erg and HB but demonstrates a high activity with N(π)-methyl HB. The possibility that the substrate is a fraction of His(methyl)_4_ in which one methyl group is bonded to N(π) and three others are distributed among N(α), N(τ), and O positions (i.d., the α-amino group is incompletely methylated) is unlikely because all related MIO-deficient lyases exhibit a strong requirement for a trimethylamino group in their substrates [20].

N(π)-Methyl HB is not commercially available, which prevented determining the conditions for inducing the corresponding lyase activity in *S. oneidensis* cells and their ability to use this substrate as a terminal electron acceptor for anaerobic growth. Nevertheless, indirect evidence suggests that the flavocytochrome *c* encoded by the *so_3300–3301* genes may convert N(π)-methyl urocanate. Specifically, the active site in the modelled structure of the flavocytochrome SO_3301 subunit is very similar to that of the urocanate reductase SO_4620 of the same bacterium (Figs 8 and S4), and its size in SO_3301 is sufficient to accommodate the additional N(π) methyl group. Furthermore, the observed differences in active site structures are fully consistent with the differences in the structures of the corresponding substrates.

It is therefore highly likely that the proteins SO_3299–SO_3301 confer to *S. oneidensis* the ability to use N(π)-methyl urocanate, formed from N(π)-methyl HB, as a terminal electron acceptor in anaerobic respiration. These proteins are thus functionally similar to the SO_3056–SO_3058 system but are tuned to a different histidine derivative. The high catalytic efficiency of SO_3299 toward N(π)-methyl HB (*k*_cat_/*K*_m_ of 4.1 × 10^6^ M^-1^· s^-1^) makes it a likely candidate for the natural substrate of SO_3299, but no data indicate the presence of this compound in nature.

Alternatively, the natural substrate of SO_3299 may be a close N(π)-methyl HB analog, similarly containing methyl groups on the N(α) and N(π) atoms. Ovothiols [38] (Fig. 2), widespread natural sulfur-containing antioxidants, are the best candidates, insofar as various species of the genus *Shewanella* encode a protein (OvoA, SO_4465 in *S. oneidensis*) for ovothiol synthesis [38]. The naturally occurring histidine derivatives ovothiols B and C contain a methyl group at the N(π) position and one or two methyl groups at the N(α) position (Fig. 2). However, it is unclear whether partial methylation of the N(α) atom is sufficient for these ovothiols to be substrates for SO_3299– SO_3301. A more promising ovothiol with three methyl groups at this position (hypothetical “ovothiol D”) has not yet been found. Thus, further research is clearly required to determine the true natural substrate for SO_3299–SO_3301-mediated respiration in *S. oneidensis*.

Gene clusters encoding putative periplasmic trimethylammonia-lyase and flavocytochrome *c* are found in many anaerobic bacteria including various *Shewanella, Ferrimonas, Sutterella, Turicimonas, Limnobaculum, Citrobacter, Hafnia, Enterobacillus, Desulfobaculum*, and *Campylobacter* species. This observation suggests that the ability to use the 2-enoates formed from HB [9] or from Erg and N(π)-methyl HB (this work) as terminal electron acceptors for the electron transport chain is widespread among many anaerobic and facultatively anaerobic bacteria.

## CRediT authorship contribution statement

Yulia Bertsova: Methodology, Investigation. Marina Serebryakova: Methodology, Investigation. Olga Godovanets: Investigation. Victor Anashkin: Methodology, Investigation. Alexander Baykov: Data curation, Visualization, Writing. Alexander Bogachev: Conceptualization, Methodology, Investigation, Writing.

## Declaration of competing interest

The authors declare that they have no financial or personal relationships with other people or organizations that could inappropriately influence or bias their work.

## Acknowledgements

MALDI MS and laser scanner facilities became available to us in the framework of the Moscow State University Development Program PNG 5.13. We are grateful to Drs. N.S. Melik-Nubarov and M.D. Mamedov for help with methylation experiments.

## Funding

This work was supported by the State assignment of Lomonosov Moscow State University.

## Data availability

All data are available from the authors at a reasonable request.

## Notes

### Competing Interest Statement

The authors have declared no competing interest.

