## Supplemental Figures S1-S6 for "Trimethylammonia-lyases of *Shewanella oneidensis* and Their Role in Anaerobic Respiration"

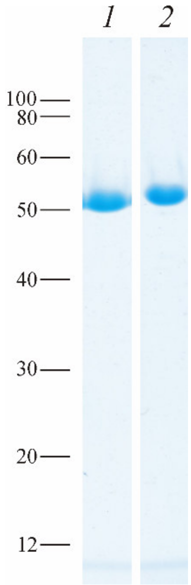


**Fig. S1.** SDS-PAGE of the isolated SO_3057 (lane *1*) and SO_3299 (lane *2*). The gel was stained with Coomassie Blue. The protein load was 4 μg per lane. The bars with numbers on the left side denote the positions and molecular masses of marker proteins.

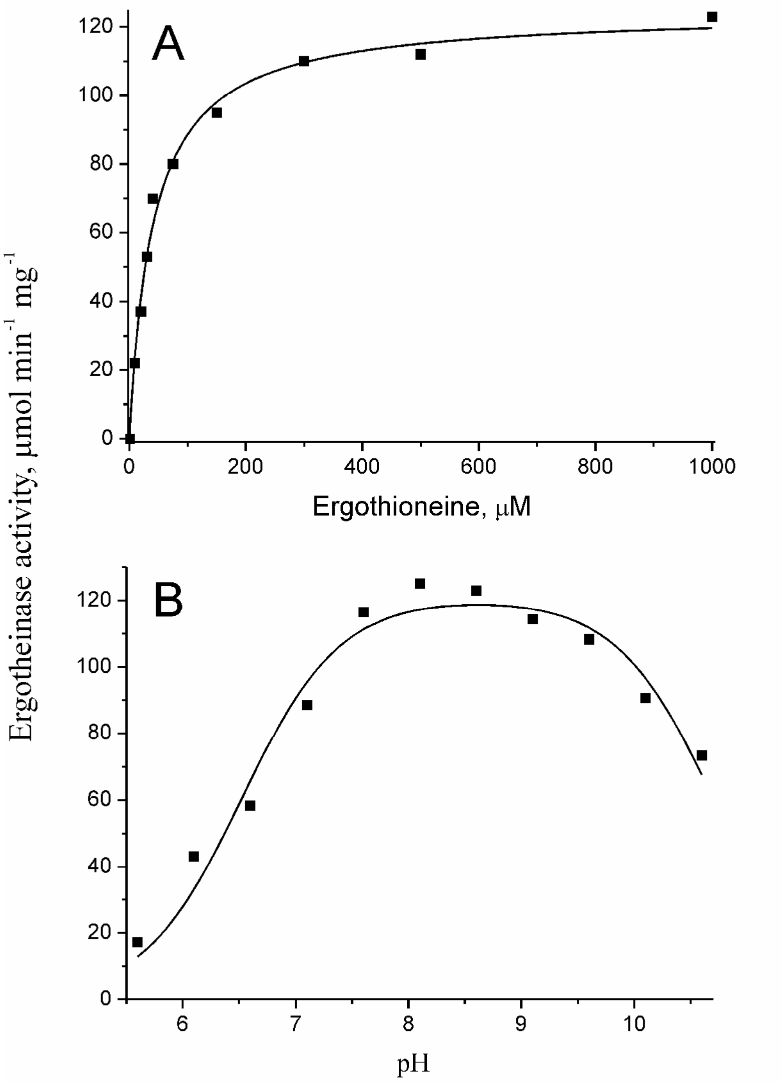


**Fig. S2.** Dependence of the SO_3057 activity on Erg concentration (A) and pH (B). The lines show the best fits of the Michaelis-Menten equation and Eq. 1, respectively.

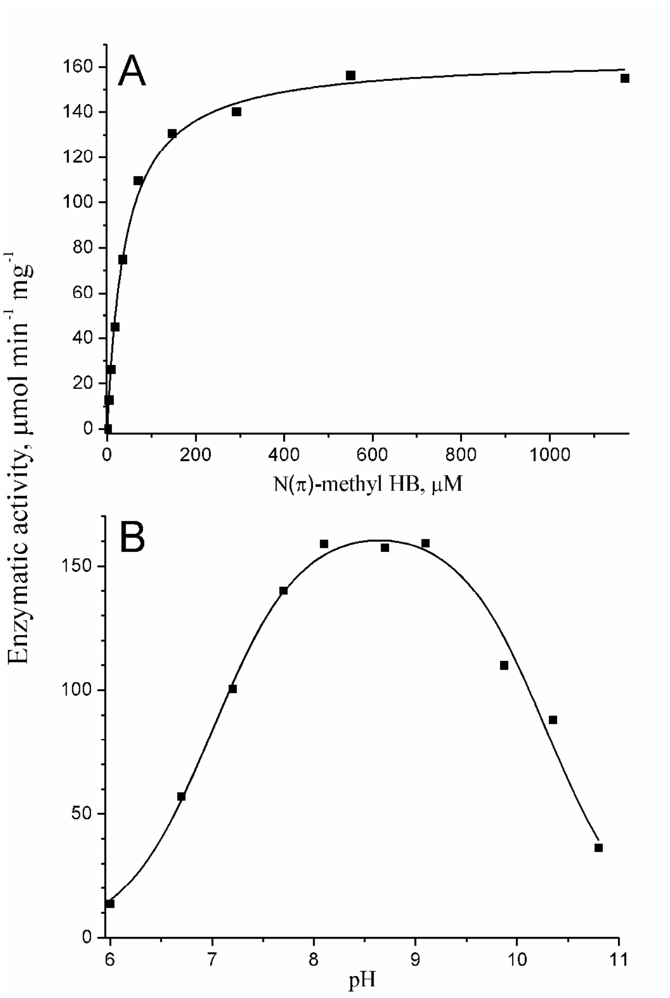


**Fig. S3.** Dependence of SO_3299 activity on N(π)-methyl HB concentration (A) and pH (B). The lines show the best fits of the Michaelis-Menten equation and Eq. 1, respectively.

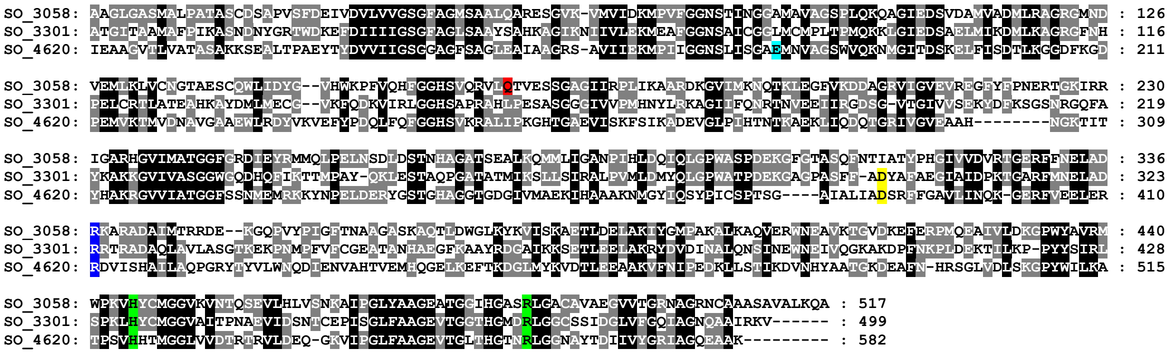

**Fig. S4.** Multiple sequence alignment of the SO_3058 and SO_3301 proteins and urocanate reductase (SO_4620) of *S. oneidensis*. The alignment was produced with ClustalW. The residues involved in the binding of urocanate carboxylic group, imidazole N(π)-atom, and imidazole N(τ)-atom are highlighted in green, turquoise, and yellow, respectively. Other colors: blue, the proton donor Arg residue participating in double bond reduction; red, the Gln residue bonded to the S-atom of thiourocanate.

```
shl_Shal_3447          : PRVLQTVQSSGAGIIRPLIKAA : 187     PWASPDEKGFGTASQFNTIATYPS : 320
spl_Spea_3368          : PRVLQTVQSSGAGIIRPLIKAA : 187     PWASPDEKGFGTASQFNTIATYPS : 320
shey_FS418_10025       : PRVLQTVESSGAGIIRPLIKAA : 187     PWASPDEKGFGTASQFNTIATYPS : 320
seur_FM038_008445      : PRVLQTVESSGAGIIRPLIKAA : 187     PWASPDEKGFGTASQFNTIATYPS : 320
sse_Ssed_0344          : PRVLQTVESSGAGIIRPLVKAA : 187     PWASPDEKGFGTASQFNTIATYPR : 320
slo_Shew_2281          : PRVLQTVQSSGAGIIRPLIKAA : 187     PWASPDEKGFGTASQFNTIATYPS : 320
saeg_K0H80_11635       : PRVLQTVQSSGAGIIRPLIKAA : 187     PWASPDEKGFGTASQFNTIATYPS : 320
shns_K0J45_11645       : PRVLQTVQSSGAGIIRPLIKAA : 187     PWASPDEKGFGTASQFNTIATYPS : 320
shao_K0H81_08270       : PRVLQTVQSSGAGIIRPLIKAA : 187     PWASPDEKGFGTASQFNTIATYPS : 320
sspa_K0I31_11905       : PRVLQTVQSSGAGIIRPLIKAA : 187     PWASPDEKGFGTASQFNTIATYPS : 320
srhs_K0I63_11830       : PRVLQTVQSSGAGIIRPLIKAA : 187     PWASPDEKGFGTASQFNTIATYPS : 320
salg_BS332_16325       : QRVLQTVESSGAGIIRPLVKAA : 187     PWASPDEKGFGTASQFNTIATYPH : 320
schk_GII14_13700       : QRVLQTVESSGAGIIRPLVKAA : 187     PWASPDEKGFGTASQFNTIATYPH : 320
scaa_TUM17_24080       : QRVLQTVESSGAGIIRPLVKAA : 187     PWASPDEKGFGTASQFNTIATYPH : 320
shej_MZ182_12220       : QRVLQTVESSGAGIIRPLINAA : 185     PWASPDEKGFGTASQFNTIATYPH : 318
shw_Sputw3181_1576     : QRVLQTVESSGAGIIRPLINAA : 185     PWASPDEKGFGTASQFNTIATYPH : 318
spc_Sputcn32_2432      : QRVLQTVESSGAGIIRPLINAA : 185     PWASPDEKGFGTASQFNTIATYPH : 318
son_SO_3058            : QRVLQTVESSGAGIIRPLIKAA : 185     PWASPDEKGFGTASQFNTIATYPH : 318
shf_CEQ32_01540        : QRVLQTVESSGAGIIRPLIKAA : 185     PWASPDEKGFGTASQFNTIATYPH : 318
sxm_MKD32_07520        : QRVLQTVESSGAGIIRPLIKAA : 185     PWASPDEKGFGTASQFNTIATYPH : 318
smay_K0H60_07765       : QRVLQTVESSGAGIIRPLIKAA : 185     PWASPDEKGFGTASQFNTIATYPH : 318
sdeo_D0436_08505       : QRVLQTVESSGAGIIRPLIKAA : 185     PWASPDEKGFGTASQFNTIATYPH : 318
shn_Shewana3_1489      : QRVLQTVESSGAGIIRPLIKAA : 185     PWASPDEKGFGTASQFNTIATYPH : 318
shel_JM642_07445       : QRVLQTVESSGAGIIRPLIKAA : 185     PWASPDEKGFGTASQFNTIATYPH : 318
shen_FJD32_008350      : QRVLQTVESSGAGIIRPLIKAA : 185     PWASPDEKGFGTASQFNTIATYPH : 318
sbj_CF168_13310        : QRVLQTVESSGAGIIRPLIKAA : 185     PWASPDEKGFGTASQFNTIATYPH : 318
fbl_Fbal_3138          : PRILQTRESSGAGIIRPLIKVA : 187     PWSSPDEKGFGTASQFNTIATYQN : 320
lri_NCTC12151_02942    : PRTLQTVESSGAGIVMPLRNSA : 195     PWSSPDEKGFGNVSQFNTIAGYPM : 328
lgb_V8L73_15360        : PRTLQTVESSGAGIVMPLRNSA : 196     PWSSPDEKGFGNVSQFNTIAGYPM : 329
prag_EKN56_16295       : PRTLQTVESSGAGIIMPMRQSA : 202     PWSSPDEKGFGNVSQFNTIAGYPM : 335
ciy_HAP28_25260        : PRTLQTVESSGAGITRPLKESA : 187     PWSSPDERGFGYVSQFNTISGFPN : 320
caf_AL524_03065        : PRTLQTVESSGAGITRPLKESA : 187     PWSSPDERGFGYVSQFNTISGFPN : 320
cir_C2U53_09755        : PRTLQTVESSGAGITRPLKESA : 187     PWSSPDERGFGYVSQFNTISGFPN : 320
cbra_A6J81_17205       : PRTLQTVESSGAGITRPLKESA : 187     PWSSPDERGFGYVSQFNTISGFPN : 320
eal_EAKF1_ch2217       : PRTLQTVESSGAGITRPLKDSA : 187     PWSSPDERGFGYVSQFNTIAGFPN : 320
ebb_F652_2006          : PRTLQTVESSGAGIVMPLKNSA : 188     PWSSPDERGFGYVSQFNTISGFPN : 321
hpar_AL518_11085       : PRTLQTVESSGAGIVMPLKNSA : 186     PWSSPDERGFGYVSQFNTISGFPN : 319
pbiz_LWC08_01865       : PRTLQTTISSGAGIVRPLRKSA : 202     PWSSPDEKGFGMVSQFNTIAGFPM : 335
sutt_SUTMEG_07030      : PRTLQTVESTGGGITRPLTESC : 179     PWCSPDERGFGLVSQFNTIAGFPM : 311
bbay_A4V04_10515       : PRIMQTVESTGGGITRPLVEAA : 178     PWSSPDERGFGLVSQFNTIAGFPK : 311
```

**Fig. S5.** Partial sequence alignment of putative bacterial thiourocanate reductases in the regions containing amino acid residues forming thiourocanate-binding site. These residues correspond to Gln168, Trp296, Phe304, Ser308, Gln309, and Thr312 of the protein SO_3058 from *S. oneidensis* (Fig. 8B) and are highlighted in purple. The bacterial flavocytochrome sequences shown were taken from the KEGG database and their genes belong to the operons additionally containing ergothionase genes. The host bacteria are as follows: *Shewanella oneidensis* (son), *S. sp.* FDAARGOS_354 (shf), *S. xiamenensis* (sxm), *S. mangrovisoli* (smay), *S. sp.* LZH-2 (shel), *S. sp.* LC6 (shen), *S. decolorationis* (sdeo), *S. sp.* ANA-3 (shn), *S. bicestrii* (sbj), *S. sp.* JNE2 (shej), *S. sp.* W3-18-1 (shw), *S. putrefaciens* CN-32 (spc), *S. carassii* (scaa), *S. algae* (salg), *S. chilikensis* (schk), *S. spartinae* (sspa), *S. halifaxensis* (shl), *S. pealeana* (spl), *S. loihica* (slo), *S. aegiceratis* (saeg), *S. alkalitolerans* (shns), *S. halotolerans* (shao), *S. rhizosphaerae* (srhs), *S. sediminis* (sse), *S. sp.* YLB-09 (shey), *S. eurypsychrophilus* (seur), *Ferrimonas balearica* (fbl), *Desulfobaculum bizertense* (pbiz), *Leminorella richardii* (lri), *L. grimontii* (lgb), *Limnobaculum zhutongyuii* (prag), *Sutterella megalosphaeroides* (sutt), Burkholderiales bacterium YL45 (bbay), *Escherichia albertii* (eal), Enterobacteriaceae bacterium bta3-1 (ebb), *Hafnia paralvei* (hpar), *Citrobacter sp.* CFNIH10 (cir), *C. sp.* Y3 (ciy), *C. amalonaticus* (caf), and *C. braakii* (cbra).
